# Wagers for work: Decomposing the costs of cognitive effort

**DOI:** 10.1101/2023.07.25.550227

**Authors:** Sarah L. Master, Clayton E. Curtis, Peter Dayan

## Abstract

Some aspects of cognition are more taxing than others. Accordingly, many people will avoid cognitively demanding tasks in favor of simpler alternatives. Which components of these tasks are costly, and how much, remains unknown. Here, we use a novel task design in which subjects request wages for completing cognitive tasks and a computational modeling procedure that decomposes their wages into the costs driving them. Using working memory as a test case, our approach revealed that gating new information into memory and protecting against interference are costly. Critically, other factors, like memory load, appeared less costly. Other key factors which may drive effort costs, such as error avoidance, had minimal influence on wage requests. Our approach is sensitive to individual differences, and could be used in psychiatric populations to understand the true underlying nature of apparent cognitive deficits.

**Author Summary:** Anyone who has tried to mentally calculate how much to tip at a restaurant knows that cognitive effort can feel aversive. Doing math in your head, like most high-level cognitive abilities, depends critically on working memory (WM). We know that WM is sometimes effortful to use, but we don’t know which aspects of WM use drive these effort costs. To address this question, we had participants request wages in exchange for performing various tasks that differed in their specific WM demands. Using computational models of their wage demands, we demonstrated that some aspects of WM are costly, such as bringing new information into memory and preventing interference. Other factors, like the amount of information in memory and attempts to avoid mistakes, were less costly. Our approach identified which specific subcomponents of WM are aversive. Future research could use these methods to test theories about how motivational problems might be masquerading as cognitive deficits in psychiatric populations.

## Introduction

Some activities (e.g., getting dinner with friends) are more enjoyable than others (e.g., calculating how to split the bill). Doing tasks which require greater cognitive effort, colloquially called “brain power,” can feel uniquely aversive, though to different degrees for different people (1–3). Indeed, despite tangible benefits, people often avoid cognitively demanding work (4,5). Such resistance suggests that we weigh the effort of mental activity, perhaps as a cost to be offset with reward.

Previous research has identified the experimental tasks which are more costly to perform by giving subjects control over which tasks they complete. Tasks which subjects demanded the most incentives to complete (5–7) or which subjects tended to avoid in favor of other tasks with equivalent rewards (4,8,9) are considered most effortful. Some costly aspects of these tasks are external, like time on task (10–12) or the complexity of the cognitive model required by the task (13–16), but other costs arise from the internal operations necessary to realize external actions. In general, cognitive resistance increases when tasks place substantial demands on working memory and cognitive control (5,17–19). However, it remains unknown which particular aspects of working memory and cognitive control may be most costly. For example, perhaps the sustained effort required during working memory maintenance is more costly than the transient effort required to inhibit a prepotent response.

Here, we decomposed simple and complex attention (i.e., detection) and working memory (N-back) tasks into putative elemental processes such as maintaining information in memory of different loads and resisting interference from task irrelevant lures. We assumed that the subjective costs of these operations are internally felt and consciously accessible, and that the total cost of completing a task is learned by experiencing these costs. We assessed these total costs using a modified auction procedure. Previous work has used such auctions to infer the subjective values of items on a menu (20,21); our modifications allowed us to infer the total effort costs associated with completing various cognitive tasks by asking subjects what a “fair wage” for task completion would be. Given the evidence that the allocation of cognitive resources is subject to a cost-benefit tradeoff (12,22–26), we hypothesized that subjects’ trial-by-trial fair wage demands would, at least to a first approximation, reflect the sum of the individual costs associated with task completion, as the amount of reward necessary to offset them. To assess the extent to which the costs we measured were related to the self-reported tendency to engage in effortful cognitive tasks, albeit varying across trials, we collected Need For Cognition scores from each of our subjects (NFC; 2).

As our subjects were likely to experience costs other than those deriving from cognitive effort, we designed our experiment to try to limit the effects of these other factors. First, to minimize the influence of time on task on fair wage ratings, we gave subjects an easy task to complete when they wished to skip a harder one. We also ensured that every task round took the same amount of time. Second, cognitively effortful tasks often also elicit errors. This may be experienced as a cost, particularly in perfectionist subjects (27,28). While we could not as easily control for error avoidance costs as for time costs, we designed our task to minimize error avoidance behavior by not giving trial-by-trial feedback, not informing subjects of their accuracy round-by-round, and not reducing their compensation unless errors became overly prevalent. We also collected subject scores on the Short Almost Perfect Scale (SAPS; 29), to assess the degree to which subjects’ fair wages were driven by the tendency to avoid making errors (i.e. perfectionism). Lastly, we included the costliness of errors alongside the costs of cognition in our computational analyses.

We found three non-zero cognitive effort costs: the cost of adding new information into working memory (WM), the cost of filtering out irrelevant information, and the cost of maintenance. More subtly, we found evidence that subjects learned the total costs of each task through task experience, and that the costs of cognition did not increase or decrease over the duration of the experiment. We found that the self-reported tendency to avoid effort was related to explicit ratings of task costliness and difficulty, as well as more implicit costs of cognition. This implies that effort avoidance may be driven both by the explicit, stable preference to avoid effort and by the implicit subjective experiences of the costs of cognition.

## Results

100 subjects completed the experiment online through Amazon Mechanical Turk. Subjects completed 32 task rounds and performed four different tasks in random order: an attentional vigilance task (1-detect), a vigilance task requiring more WM maintenance (3-detect), and the 1- and 2-back WM task (18,30). Before each task round began, subjects were shown the task they were to complete (an associated fractal), and were able to request a fair wage for that round of that task. A Becker-Degroot-Marschak (BDM) auction mechanism then determined whether they completed 15 trials of the task they rated for the wage they requested, or 15 trials of the default, non-demanding task (the 1-detect) for a lower wage. We analyzed their performance and fair wage ratings across tasks. We used computational modeling to examine how fair wages were influenced by the putative cognitive operations used to complete the previous task rounds, like WM maintenance or updating. We also related fair wage ratings to previous task behavior, including the number and types of errors they made.

### Model-Agnostic Results

There was a main effect of task identity on accuracy (Figure 2A; F = 44.1; p <0.0001), mean reaction time (RT; F = 31.5, p <0.0001), and difficulty rating (F = 26; p <0.0001). Post-hoc comparisons confirmed that subjects had lower accuracy, higher RTs, and higher difficulty ratings on the 2-back task than on all of the other tasks (Table 1; Accuracy: 2-back versus 1-detect p <0.001; 2-back versus 1-back p <0.0001; 2-back versus 3-detect p <0.0001; Mean RT: 2-back versus 1-detect p < 0.001; 2-back versus 1-back p <0.0001; 2-back versus 3-detect p <0.0001; Difficulty ratings: 2-back versus 1-detect p <0.001; 2-back versus 1-back p <0.0001; 2-back versus 3-detect p <0.0001). Accuracy was highest on the 1-detect when compared with all the other tasks (all p’s <0.0001). Mean RTs on the 1-detect were lower than on the 1-back (p <0.0001), and 2-back (p <0.0001), but not on the 3-detect (p >0.05). Difficulty ratings were also lowest on the 1-detect compared to the 1-back (p <0.0001), 3-detect (p <0.0001), and 2-back (p <0.0001). Mean accuracy was lower and mean RT was higher on the 1-back than on the 3-detect (p <0.001; p <0.0001). The mean difficulty rating was no different between the 1-back and 3-detect (p >0.05).

**Figure 1.**
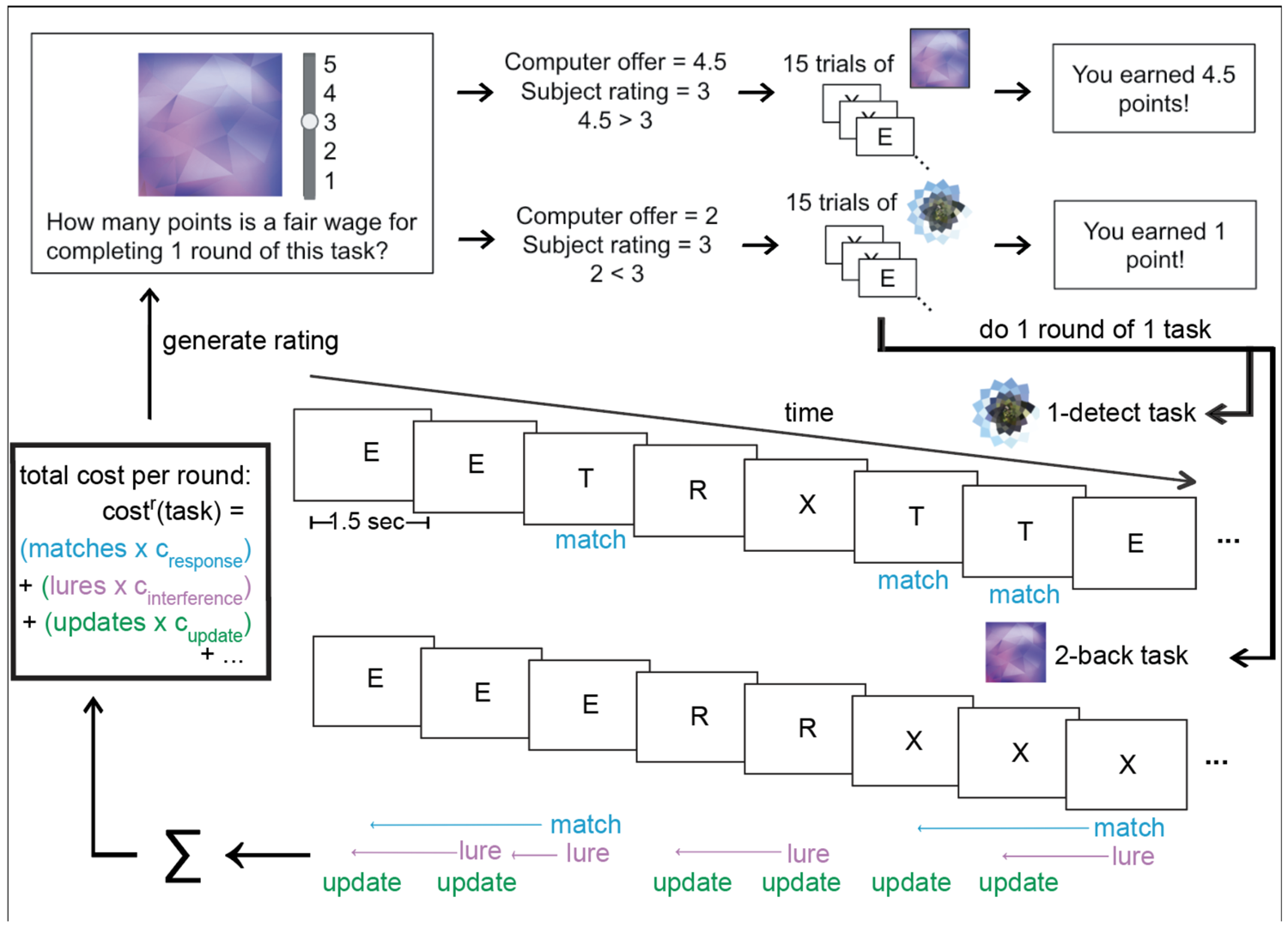
The behavioral paradigm & computational modeling approach. Before each round of the experiment, subjects were shown an image which was associated with one of three possible tasks. They then indicated the wages (in points) that they would like to receive for completing 1 round of that task. If their fair wage rating was below a random computer offer, then they would complete that task and receive the computer’s offer. If their fair wage was above a random computer offer, then they would complete a different, easier task instead. We employed this inversion of the Becker-Degroot-Marschak auction procedure to incentivize subjects to be truthful in their fair wage ratings.

**Figure 2.**
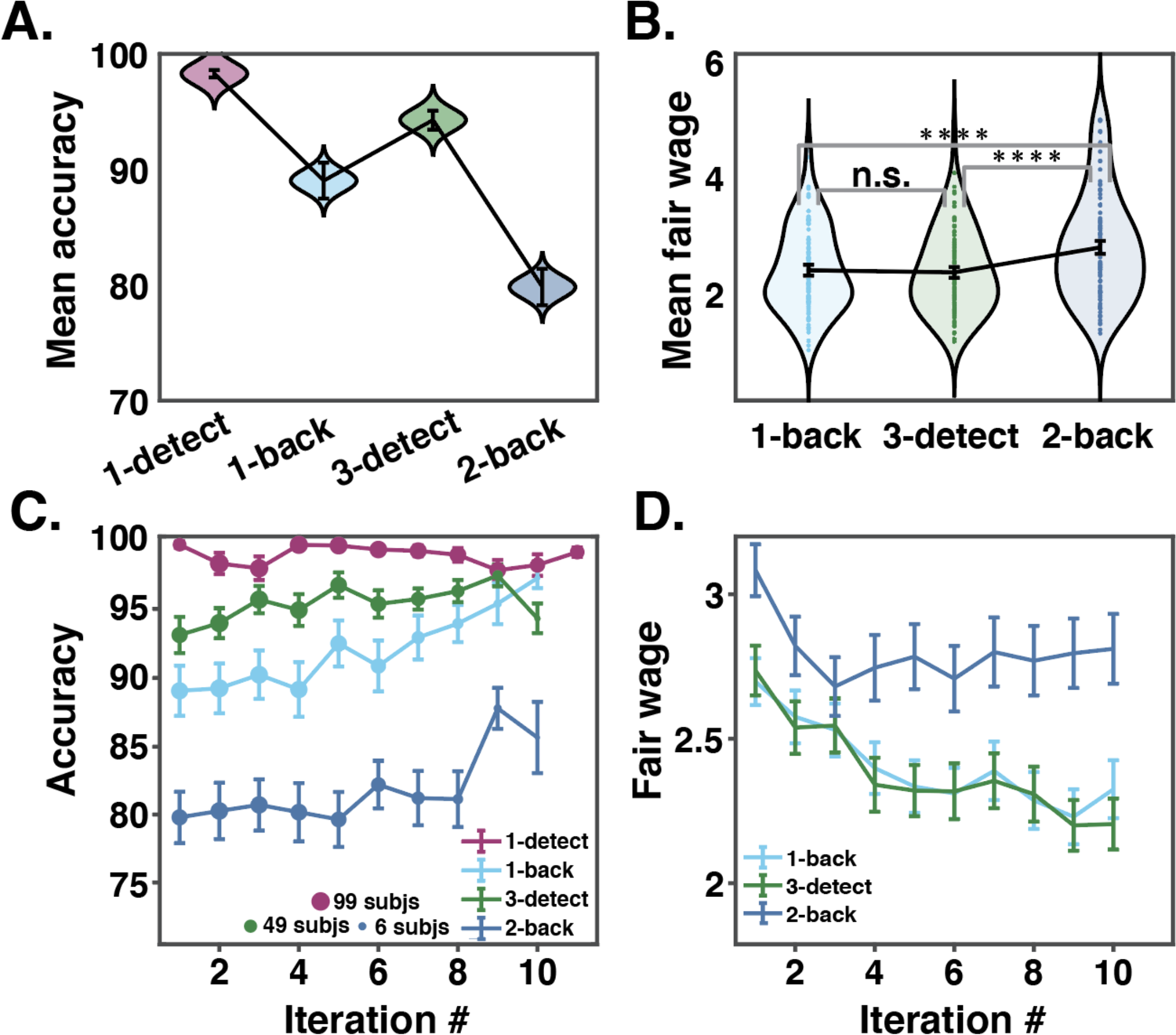
Model-agnostic behavioral results. **A.** Distributions of mean accuracies across all subjects for the default task (1-detect), and the three rated tasks (1-back, 3-detect, and 2-back). The black bars depict the means and standard errors of the mean (SEMs) of each distribution. The distribution of all subjects’ mean accuracies was plotted using a Gaussian kernel via violin.m. All mean accuracies for each task were significantly different from each other (all p’s < 0.001). **B.** Distributions of mean fair wages across all subjects for the three rated tasks. The lowest possible rating was 1, and the highest possible rating was 5. The black bars depict the means and SEMs of each distribution. The distribution of ratings was plotted using violin.m. **** indicates significance at the p < 0.0001 level. **C.** Mean accuracy across all subjects on each iteration of each task. Due to the stochasticity inherent to the BDM auction procedure, individual subjects completed the 1-back, 3-detect, and 2-back tasks a variable number of times, but a maximum of 11 times each. The relative number of subjects who completed each iteration is depicted by the size of the dot plotted at the mean. Error bars are plotted with standard error of the mean. A two-way ANOVA of task and task iteration revealed a main effect of task identity (F = 15, p < 0.0001) but no effect of task iteration (F = 1.3, p > 0.05). Thus mean accuracy was different across tasks but did not change with task experience. **D.** Mean fair wage rating by rating number, where the maximum is 11 ratings of one task. A 2-way ANOVA on BDM ratings showed a main effect of task identity (Table 1; F = 33; p < 0.0001) and a main effect of task iteration (Figure 1; F = 21; p < 0.0001).

**Table 1:** Mean accuracy, reaction time (RT) in milliseconds, and difficulty ratings across all subjects for the default task, the 1-detect, and for the three rated tasks, the 1-back, 3-detect, and 2-back tasks. The maximum RT was 1500 milliseconds. The minimum fair wage and difficulty rating was a 1, and the maximum was a 5.

| Group means | **1-detect** | **1-back** | **3-detect** | **2-back** |
| --- | --- | --- | --- | --- |
| **Percent accuracy** | 98.29 | 89.03 | 94.28 | 79.83 |
| **RT (msec)** | 548.33 | 610.97 | 532.59 | 717.37 |
| **Difficulty rating** | 1.90 | 2.44 | 2.40 | 3.32 |
| **Fair wage** | NA | 2.41 | 2.37 | 2.80 |

Subjects rated only the 1-back, 2-back, and 3-detect tasks, as the 1-detect task was the default task. A 2-way ANOVA on fair wage ratings showed a main effect of task identity (Figure 2B; Table 1; F = 29.7, p < 0.0001) and a main effect of task iteration (Figure 2D; Supplementary Figure 1; F = 5.2, p < 0.0001). Subjects’ mean fair wage ratings on the 2-back task were significantly higher than for the 1-back (p < 0.0001). Comparing fair wage ratings for the 1- and 2-back allows us to directly measure the costs of maintaining one more item in working memory, though the 1- and 2-back tasks also differ in the degree of interference present in WM and the number of errors made. Mean fair wage ratings on the 2-back were also higher than on the 3-detect (p < 0.0001). Comparing fair wages from the 2-back and 3-detect, which both require the maintenance of 2 items, allows us to measure the cost of interfering stimuli in WM storage or the increased errors made on the 2-back task. Fair wages were not significantly different between the 1-back and the 3-detect (p > 0.1). Though the 1-back and 3-detect differ in their load on WM, subjects tended to rate them equivalently. These results suggest that increasing WM interference may be more subjectively costly than increasing WM load. We investigate further in our model-based analyses below.

While accuracy was significantly lower and fair wages significantly higher for the 2-back task, there was no relationship between mean 2-back accuracy and mean fair wage on the 2-back across subjects (r = −0.08, p > 0.1). There was also no relationship between mean accuracy and fair wage on the 3-detect task (r = −0.16, p > 0.1). However, there was a significant relationship of mean 1-back accuracy and mean fair wage (r = −0.36, p < 0.01). We further assess the influence of errors on fair wages below by using computational modeling (Model-based results).

Task accuracy was broadly stable across task iterations (Figure 2C; main effect of task iteration F = 1.3, p > 0.05). This indicates that performance did not improve with task experience. Across all subjects there was also no relationship between round number (out of 32) and mean task accuracy or mean RT (Pearson r = −0.02, p > 0.1; Pearson r = 0.003, p > 0.1). This is likely because subjects trained to 80% accuracy during practice and were already at their maximum performance levels by the start of the main task.

Fair wage ratings seem to decrease with task iteration (F = 5.2, p < 0.0001; Figure 2D), but potentially as a byproduct of the experimental design. That is, subjects who asked for lower wages completed the non-default tasks a higher number of times; therefore the lower mean fair wage on later task iterations may primarily come from subjects who had lower fair wage ratings overall (Supplementary Figure 2). Another possibility is that subjects ask for lower fair wages over time because they find that the tasks become less effortful with practice. If that were the case, then you might expect their accuracy to improve over the course of the experiment. However, the ANOVA on task accuracy by task iteration reported above found no main effect of task iteration. We investigated this further by averaging fair wages over each subject’s first and last half of task completions, and comparing them via t-test to see whether their wage requests changed as their task experience increased. We did the same analysis for task accuracy. There was a significant decrease of fair wage ratings from the first to the second half of task completions for the 1-back task (p < 0.01) and 3-detect task (p < 0.01). There was no change in fair wage ratings across the first and second halves of experience with the 2-back task (p > 0.05). There was no change in accuracy in the first and second halves of task completions on the 1-back task (p > 0.05), 3-detect task (p > 0.05), or 2-back task (p > 0.05). Taken together, these results suggest that any decrease of fair wage ratings over task iterations stems from the experimental design, and not from learning or practice effects. We investigate this further with computational modeling below.

### Analysis of Self-Report Measures

We ran regressions on task behavior with linear and quadratic NFC and SAPS terms, using a model selection procedure which trimmed each regression down to an intercept term, and the self-report terms which were necessary for model significance (p < 0.05). NFC scores were linearly and quadratically related to mean 3-detect accuracy (β = −11.59, β = 1.82). NFC was quadratically related to difficulty ratings for the 1-detect (β = −0.06). SAPS scores were linearly and quadratically related to mean 1-back accuracy (linear β = 12.94, quadratic β = −1.60), mean 3-detect accuracy (linear β = 7.58, quadratic β = −0.87), and difficulty ratings for the 2-back task (linear β = −1.15, quadratic β = −0.13). SAPS scores were also quadratically related to 2-back accuracy (β = −1.11). Neither NFC nor SAPS score were linearly or quadratically related to mean RTs.

We ran the same regression analysis on mean fair wage ratings, collapsed across all tasks. There was a significant quadratic relationship of NFC and mean fair wage ratings (β = −0.03). We split subjects up into self-report tertiles to further investigate the significant quadratic relationships between task and self-report variables. The tertile split resulted in 25 low, 37 mid, and 37 high NFC subjects, and 34 low, 37 mid, and 28 high SAPS subjects. Post-hoc t-tests confirmed that the significant quadratic effect of NFC is driven by the difference in mean fair wages between the high and mid NFC subjects. Mid NFC subjects had higher fair wage ratings than high NFC subjects (p < 0.01; Supplementary Figure 3). However, there were no differences between the low and high NFC groups (p > 0.05), or the low and mid NFC groups (p > 0.05). We supposed that high NFC subjects would ask for the lowest fair wages, but we did not find such a pattern in explicit fair wage ratings. We next investigated how NFC was related to the implicit costs of cognition captured by our computational model.

### Model-based results

Based on the model-agnostic results, we designed and tested a series of computational models to isolate the costs of distinct cognitive processes from fair wage ratings. These models allowed us to test the hypothesis that subjects have some internal awareness of the costs of certain cognitive operations, and to estimate the magnitude of these costs. We also measured the costs associated with all types of behavioral responses, including making errors. In doing so, we assessed whether fair wage ratings also captured costs stemming from physical effort (making key presses) or error avoidance, which are not cognitive process costs but are still potential modifiers of fair wages. Error avoidance in particular could explain, to some extent, effort avoidance in behavior; we fit error costs separately to assess this possibility (28).

We fit subjects’ behavior with a series of models using the Computational Behavioral Modeling (CBM) toolbox (31). All models included a noise parameter (σ), and an initial rating parameter for each task (*init_i_*) as free parameters. One class of models assumed that the cost parameters were fixed across trials, but that the subjects learned about the total cost they experienced with a learning rate (ɑ). A separate class of models assumed that subjects’ demands reflected the cost just on the previous iteration of the task, but that the cost parameters changed linearly with trial number at a rate given by parameter (δ). Within these model classes, we tested several combinations of cost parameters. The maintenance cost (c_maintenance_) captured the effect of maintaining more information in WM. The interference cost (c_interference_) captured the effect of “lure” trials in the 2-back task. The update cost (c_update_) captured the effect of updating WM with new information. The response cost parameter (c_response_) captured the influence of perceived matches (button presses) on subsequent BDM ratings. The miss cost (c_miss_) captured the effect of missed matches. The false alarm cost (c_fa_) captured the effect of making responses when there was no match.

The models with the highest frequencies in our subject pool included learning rate ɑ, rating noise σ, three initial rating parameters (one per task), update costs, interference costs, and maintenance costs (Figure 3A). Two subjects were best fit by a model including the cost changing parameter δ and a fixed learning rate ɑ = 1 but most subjects’ (98/100) experienced costs of cognition were stable across 32 task rounds. Most changes in fair wage ratings were likely driven by cost learning (α), differences in the cognitive operations required in different task rounds, or reporting noise (σ).

**Figure 3.**
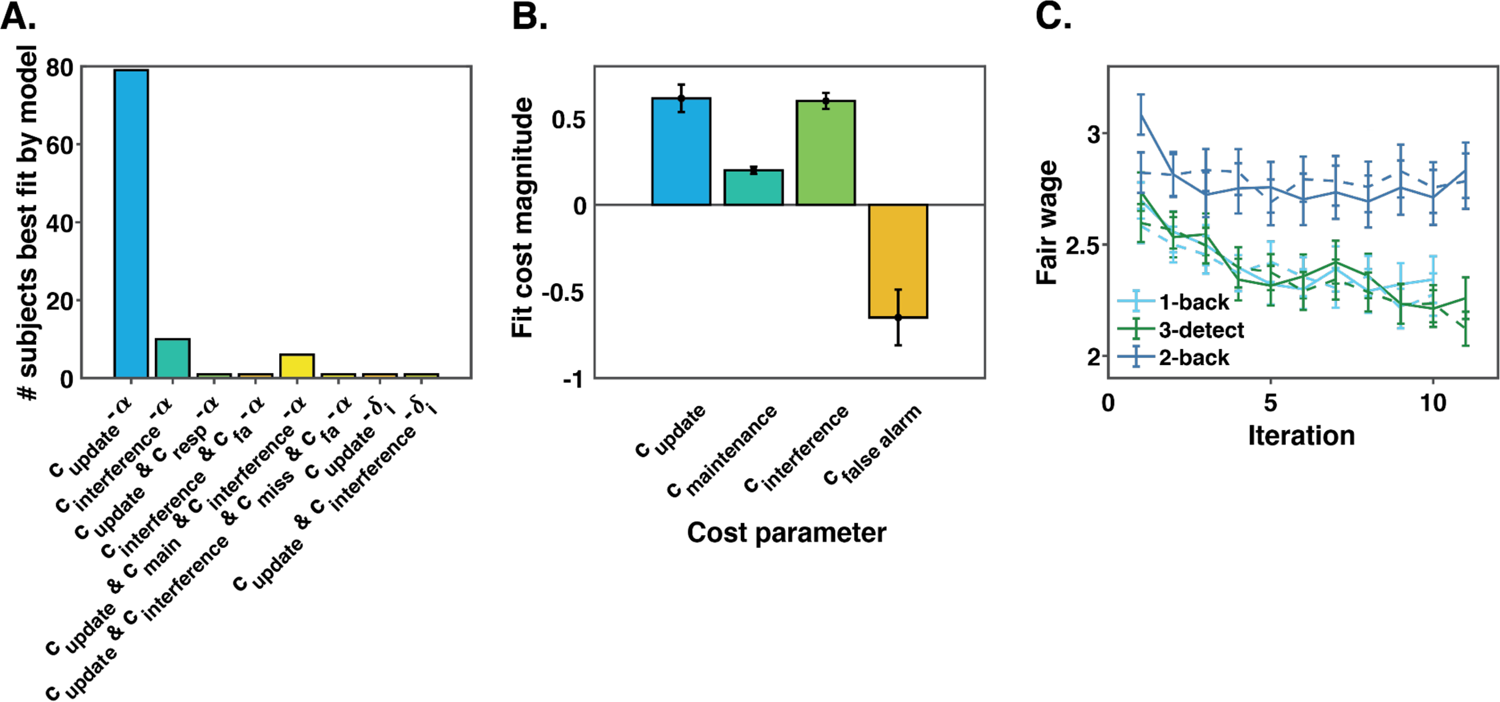
Computational modeling results. **A.** The number of subjects best fit by each model with a non-zero model frequency. Of the 84 computational models fit to subjects’ fair wages, the winning models were alpha cost-learning models containing update costs (c_update_), interference costs (c_interference_), and maintenance costs (c_maintenance_), and false alarm costs (c_fa_), in various combinations. The model with the highest model frequency was the model including update costs alone. **B.** The mean of the posterior distribution of each cost parameter from the models that best fit at least 1 subject’s fair wages. These posterior distributions were calculated by combining inferred parameter distributions across subjects and across models. Inference was performed over joint 4D distributions to capture co-variance between update, interference, maintenance, and false alarm costs. For plotting purposes we summed over the three irrelevant dimensions for each parameter to construct its marginal distribution, and then calculated the means and variances of the marginals. Error bars reflect the hierarchical standard error of the mean; they were calculated not with the square root of the total number of subjects in the denominator, but with the square root of the number of subjects’ data explained by models containing that parameter. Note that the error bars describe the spread of the marginal parameter distributions, not variance in the fitting process, and thus are not suitable for estimating the statistical significance of the effects plotted. **C.** Real (solid lines) versus simulated (dashed lines) fair wage ratings on each rating iteration for each task. Data simulated using each subjects’ best model faithfully reproduces real subject data (r^2^ = 0.51).

The model with the highest model frequency included only update costs, and was the winning model overall with a protected exceedance probability of >0.99 and a model frequency of 78.1%. The second most frequent model included only interference costs and had a model frequency of 10.3%. The third most frequent model included update, interference, and maintenance costs, and had a model frequency of 6.3%. The remaining five recovered models contained the rest of the cost components (including false alarm, miss, and response costs) in various combinations and accounted for the last 5.3% of model frequency (Figure 3A). They also contained two models with δ cost-changing parameters instead of ɑ cost-learning parameters.

Although most of our subjects were best fit by the winning model, one quarter of our subjects were best fit by other models. Subject fair wages were better fit by simulating data for each subject using their best-fitting model (mean r^2^ = 0.516; Figure 3C; Supplementary Figure 3), than by simulating data for all subjects with just the winning model (mean r^2^ = 0.466). In addition, 10 subjects’ data were best explained by models containing multiple costs of cognition. Thus, subjects’ fair wages were influenced by more than just update costs.

There was scant evidence that button presses or errors were costly, as all models including response, false alarm, or miss cost parameters had a total model frequency less than 3%. Models including response and miss costs each accounted for model frequencies less than 1%, so these costs are not explored further below.

The mean update cost was 0.615 (Figure 3B), making it the highest magnitude cost parameter. The next highest mean parameter value was the interference cost, at 0.60, followed by the maintenance cost at 0.2, and the false alarm cost, at −0.65. Despite the near equivalence of the mean update and interference costs, lures in WM were much less frequent than updates to WM. Because of this, subjects lost more monetary bonuses due to the avoidance of update costs, resulting in their forfeiting an average of 0.87 cents extra per round. They were willing to forfeit 0.26 cents and 0.38 cents per round to avoid maintenance and interference costs, respectively. While subjects did not know the exact mapping between BDM points and the monetary bonus at the conclusion of the experiment (1 point = 1 cent), this speaks to the true costliness of each component process, in terms of the overall monetary amounts subjects forfeited.

As we hypothesized, the mean difference between fair wage ratings on the 2-back and 3-detect tasks was predicted by the magnitude of the interference costs (r = 0.42, p <0.0001). The mean difference between ratings on the 2-back and 1-back was predicted by the magnitude of the maintenance costs (r = 0.41, p <0.0001). These correlations confirm that the tasks differ in their subjective costliness at least partially because of the differences in WM operations required by them.

We tested whether any self-report measures of effort avoidance or perfectionism related to fit cost parameters. Specifically, we wondered whether the need for cognition (NFC) or perfectionism (Short Almost Perfect Scale; SAPS) scales were predictive of any cost parameter values. For simplicity, we analyzed just parameter values from subjects best fit by the winning (update costs) model (N = 79). We ran a regression including both linear and quadratic terms for the effect of NFC and SAPS scores on fit update cost parameters from the winning model. We found no significant linear (β = 0.218, p >0.1) or quadratic (β = −0.044, p >0.1) relationship between update cost and NFC. There was also no significant linear (β = −0.210, p >0.1) or quadratic (β = 0.027, p >0.1) relationship between update cost and SAPS score. NFC and SAPS scores were well-sampled across our sample of 100 subjects (Supplementary Figure 2).

We then examined whether there were parameter differences across NFC and SAPS tertiles. Within the subjects best fit by the winning model, high NFC subjects had significantly lower update costs than both low (p < 0.05) and mid-NFC subjects (p < 0.05). There were also differences in initial fair wage ratings across NFC groups (Figure 4), the general pattern being that mid NFC subjects asked for the highest initial fair wages. High NFC subjects had significantly lower initial fair wage ratings than mid NFC subjects for all three tasks (1-back p < 0.01; 2-back p < 0.01; 3-detect p < 0.05). There were no significant differences between low and high NFC subjects’ initial rating parameters. Mid NFC subjects had higher initial ratings for the 2-back task than low NFC subjects (p < 0.05). Mid NFC subjects had higher variance (σ) around their fair wage ratings than high NFC subjects (p < 0.01) and low NFC subjects (p < 0.05). There were no significant differences in learning rates between subjects split into NFC tertiles (p’s > 0.1). Taken together, these results suggest that both explicit reports about task costliness (initial fair wage ratings for each task), and more implicit experiences of the costs of cognitive operations (update costs) change with individual differences in NFC across subjects. There were no significant differences in cost parameter magnitudes between subjects split into SAPS tertiles.

**Figure 4.**
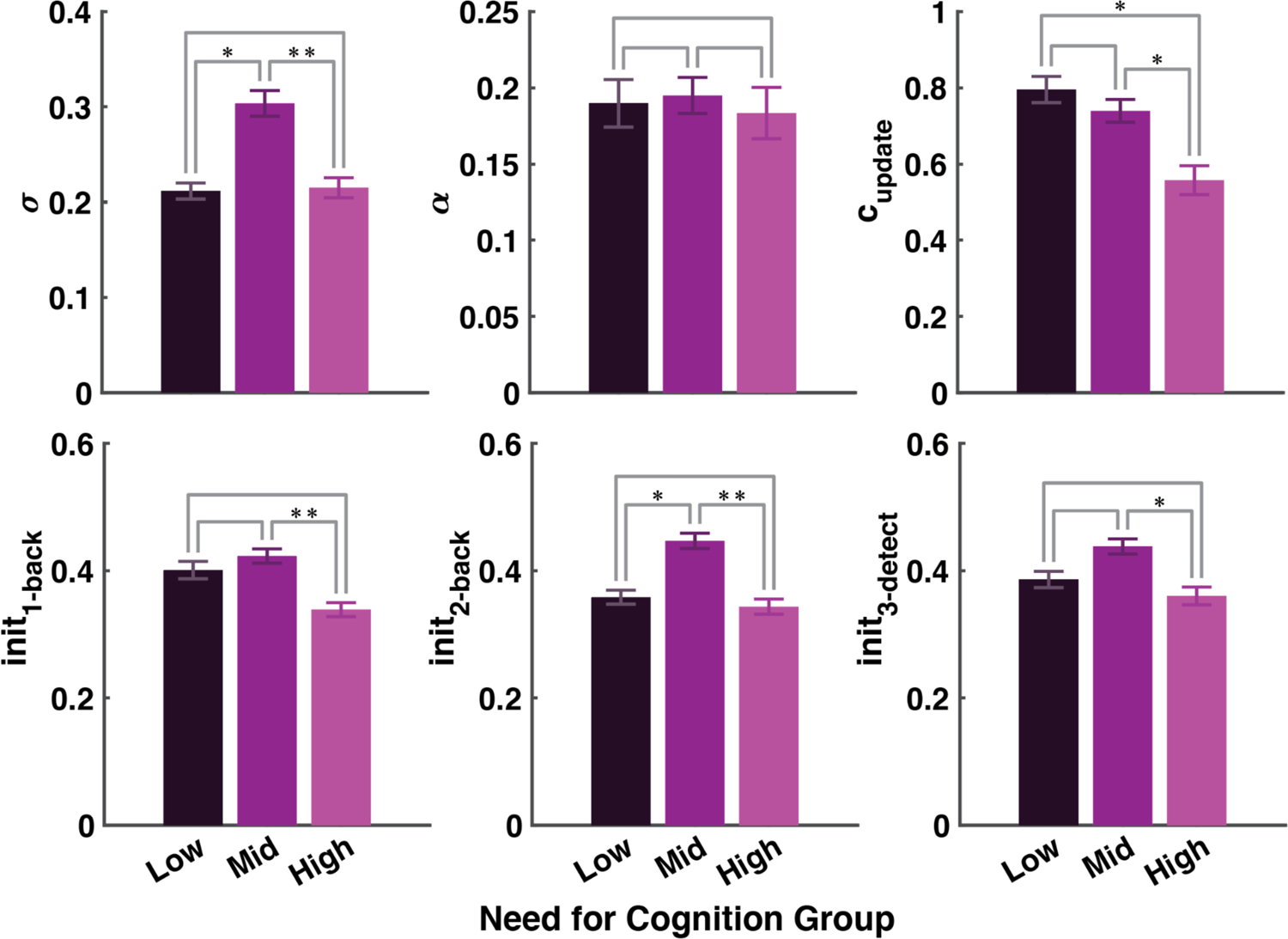
Winning model parameter values by Need for Cognition (NFC) Group. Mean parameter magnitudes from the winning 6-parameter update cost model. σ is the standard deviation parameter which dictates how noisy each subject’s fair wage ratings are, on average. ɑ is the subject-specific task cost-learning rate. The update cost is the magnitude of the influence of WM updates on each subject’s fair wage ratings. The init parameters dictate each subject’s initial fair wage for each task. Subjects were split into NFC tertiles resulting in low (N = 25), mid (N = 37), and high (N = 37) NFC groups. Fit parameter values were then averaged within-group to produce each bar. Error bars are standard error of the mean. * indicates significant difference as assessed with a t-test at p < 0.05 level. ** p < 0.01

## Discussion

Deploying working memory or paying attention can feel costly (32,33). In this work, we quantified the subjective costs of the cognitive operations demanded by commonly studied working memory and attention tasks, in a way sensitive to both the dynamics of cognitive effort exertion and individual differences in effort avoidance. Using a novel experimental paradigm which leverages an inverted Becker-Degroot-Marschak auction procedure (20), we obtained subject ratings of the total cost of completing a working memory or attention task, one round at a time. We then used a computational model to decompose these ratings into the costs of the individual cognitive operations putatively used during that round, as well as aspects of subject behavior, like errors. Our computational models quantify the subjective costs of individual cognitive operations and allow us to test several hypotheses about how cognitive effort costs may change with time or task experience.

We found evidence that updating WM, interference from within WM storage, and WM load are subjectively costly. Most subjects tracked a single cost. The largest percentage of subjects tracked just update costs, and the next highest proportion tracked just interference costs. Although effortful cognition can be rewarding (3,34), we find that the costs, not the intrinsic rewards, of cognitive effort drove fair wages. Updating WM cost the most. Subjects forfeit on average 0.87 cents extra per round as a result of avoiding frequent WM updating demands. Interference costs (lure stimuli inside of WM) were similarly high, but because lures were somewhat infrequent, rating them highly (and thereby avoiding them) led subjects to lose less money per round. The third highest cost was that of maintaining more information in WM.

Increasing WM load (the N in N-back) has often been assumed to be the primary driver of increases in subjective difficulty. However, we show that WM load was only minimally costly and that updating and interference had a greater influence on subjective cognitive effort. Lure stimuli in WM storage demand an accurate maintenance of both stimulus identity and stimulus order.

The interference cost captures the confusability of stimuli in WM storage and the cost of disambiguating them by their temporal order. WM updating is similarly complex, as information must be gated in, gated out, and temporally re-ordered. WM updating has been compared to switching between WM attractor states, which could be an energetically costly process (35,36). Perhaps, the magnitude of the update cost parameter captures the complexity or energetic costs associated with this operation.

We find that subjects quickly learned the costs of completing each task through internal cost feedback signals, then exhibited stable fair wage ratings. Our models provided two surprising new insights into how the costs of cognition may figure into deciding between several paths of action. First, only 10 subjects were best fit by models which contain multiple cost parameters. Tracking multiple costs of cognition may be in itself costly, so subjects may have selected just one cost component to base their fair wage ratings on to minimize overall experimental demands, consciously or otherwise. Second and seemingly at odds with previous work (37–39), we found no evidence that fatigue impacted fair wage ratings as cost parameters did not increase or decrease over rounds. However, cognitive fatigue may only emerge after longer durations of cognitive work (40).

Our task design directly controlled for one possible confound of the costs of cognitive effort, time on task (10–12). Another key confound in cognitive effort avoidance work is error avoidance (27,28), which is harder to directly control for, as tasks which are cognitively effortful often also elicit more errors. Instead, we measured several potential markers of error avoidance and found that it had minimal influence over subjects’ fair wage ratings. First, there was no relationship between round-by-round accuracy and fair wage ratings in two out of three tasks. Second, highly perfectionistic subjects, as measured by the Short Almost Perfect Scale (29), did not have higher fair wages overall, though they would be expected to have been particularly error avoidant. Third, while 2/100 subjects’ fair wage ratings were responsive to false alarm errors, the fit cost of making false alarms was of the smallest magnitude, and in fact, numerically negative (Figure 3B). No subject was affected by the cost of making omission errors (misses). These results suggest that while error avoidance is a small factor in the overall costs of cognitive effort, it is not the most important component driving them.

The Need for Cognition (NFC) scale measures the self-reported tendency to engage in challenging cognitive work (2). Our task and modeling approach are sensitive to self-report NFC, as the cost of updating WM is lowest in subjects with high NFC. This establishes that our paradigm is sensitive to individual differences, and validates that what we measure with it is indeed related to the trait tendency to avoid cognitive effort. Interestingly, self-report NFC scores exhibited an inverted U-shaped relationship with initial task ratings, where mid NFC subjects requested the highest fair wages on the 2-back task. This suggests that NFC interacts differently with more explicit task ratings versus the more implicit costs of cognition, and warrants further investigation. Though one would suspect that high NFC subjects would provide the lowest explicit fair wage ratings, their fair wage ratings were not significantly different from low NFC subjects’ ratings. Instead, they differed in how their wages responded to the dynamic costs of cognition (WM update costs). This suggests a dissociation of explicit self-report measures and task behavior, but an association between explicit self-reports and the implicit costs of cognition measured through computational modeling.

One limitation of our task design was the high degree of correlation between cost components, which may have impacted cost parameter recovery during model fitting. While maintenance demands were constant across the 2-back and 3-detect tasks, the 2-back was the only task which required subjects to filter out interference from lures stored in WM. In addition, as the 2-back was the most difficult task, it was associated with the most errors. Thus the total cost components increased from the 1-back to the 2-back, and to some extent from the 3-detect to the 2-back. This resulted in high correlations between cost components within subjects. Despite this consequence of the experimental design, there remained a high degree of fidelity in parameter recovery (Supplementary Figure 4), and a low degree of tradeoff between fit parameter values (Supplementary Results). It remains an open question as to what extent these cognitive operations (i.e. WM updating, resistance to interference, and maintenance) depend on overlapping or independent mechanisms, and indeed whether the costs of these operations are related.

This work directly quantifies the costs associated with the cognitive operations required in working memory and attention tasks, not just how subjects avoid or approach each task. The N-back, a classic WM task, is useful in the study of working memory because it requires the use of many diverse WM operations (30). Here, we reveal that the N-back’s strength may also be its weakness, in that the number of WM operations required to complete it is also what makes it so aversive (32).

There are many avenues for future work using this experimental and modeling approach. Here, we adopted one specific process model to decompose each round of each task into the component cognitive operations necessary to complete it, though there are many possible models to use. The use of a different process model could have resulted in a different cost component structure. Slight modifications to the tasks could also have given rise to different cost magnitudes. For example, if we had provided explicit trial-by-trial feedback, we may have observed higher error costs.

These results have potential implications for treating cognitive dysfunction in psychiatric disorders. For one, the N-back task may not be suitable for use as a benchmark for WM ability in psychiatric populations, as many have comorbid cognitive and motivational deficits. Dopaminergic cortico-striatal loops, which are highly sensitive to reward, are thought to be a driver of WM performance (41–43). Our novel paradigm may be clinically useful, as cognitive dysfunction could be partially treated by comparing the costs of cognition across groups, then offsetting those costs with rewards (44–46).

In summary, along with a novel experimental approach in which subjects request wages for completing one round of one task, we implemented a modeling procedure that decomposes their wages into the costs driving them. We found that updating WM, interference among items in WM, and WM load are costly, independent of any error, time, or fatigue costs. This suggests that certain cognitive operations are inherently costly to perform, in alignment with the idea that human cognition is subject to cost-benefit analyses which can result in the use of less costly, less effective cognitive strategies (47). Surprisingly, the highest subjective cost of N-back performance was not WM load, but WM updating. We find a direct relationship between self-report individual differences in cognitive effort avoidance and the implicit costs associated with specific WM operations. That our task captures these individual differences, where others have not (48), suggests it could be implemented to capture other individual differences, perhaps in psychiatric or developmental populations.

## Methods

100 subjects (35 female, 14 unspecified sex, mean(std) age: 39(12), 11 unspecified age) completed our online task in full. 281 unique workers opened our experiment on Amazon Mechanical Turk (AMT). Of the 270 subjects who consented to participate, 218 of them made it through the practice blocks, 142 successfully finished the quiz, 125 made it to the 16th block of the experiment, and then 100 completed the experiment in its entirety. Our final sample, which we analyze below, consisted of these 100 subjects who finished the experiment. We did not include any data from any of the subjects who did not finish the experiment in our analyses. Given the strict accuracy and attention cutoffs we imposed, and the overall length of our task (mean(median) total time on task: 37(36) minutes) versus the typical length of tasks on AMT (one study reported that the mean time spent on submitted HITs was less than 2 minutes (49)), we considered a 37% completion rate to be acceptable.

Subjects were asked to complete 32 task rounds, alternating between 4 different tasks: a 1-detect task (oddball detection), a 1-back task, a 3-detect task (detect 3 of the same stimulus in order), and a 2-back task. We chose these four tasks because they rely on many of the same cognitive processes, whilst also differing in important ways in the operations they require from those processes.

### Experimental procedure

In a novel experimental paradigm, we leveraged the Becker-Degroot-Marschak (BDM) auction procedure to measure the evolving subjective value of choice options (20). The experiment was coded using a pre-built Javascript framework for online Psychology experiments (JsPsych; 50) and custom Javascript functions. Subjects were introduced to 4 tasks, each of which was associated with a fractal image (a “task label”; see Figure 1): the 1- and 2-back working memory tasks, and two types of attentional vigilance task, which we refer to as the 1-detect (the default task) and 3-detect (4,5,30,51). Subjects completed a total of 32 rounds, using the BDM procedure before each round to report the wages they considered fair for performing the particular non-default task that was offered instead of the default 1-detect task.

In all tasks, subjects saw a sequence of 15 letters, one after the other. Subjects had to respond to the letters that matched a rule by pressing the “K” key on the keyboard. Stimuli remained on the screen for 1.5s; any response had to be made before they disappeared. If a subject responded late to a match, that trial was marked incorrect. The inter-stimulus interval was 300ms. Time on task was standardized such that the time spent on each task could not influence subjective effort cost differences across tasks; each task round took approximately 24 seconds.

The 1-detect task was the default task, intended to involve minimal effort. Subjects had to respond only if they saw a “T” on screen. In the 3-detect task, subjects had to respond when any letter was presented 3 trials in a row. In the 1- and 2-back tasks, subjects had to respond when the letter on screen matched the one displayed 1 or 2 trials back, respectively. Letter sequences were standardized such that subjects were required to respond to 3 to 5 matches per round, regardless of task identity. We chose to run these four tasks because they involved similar cognitive processes, but differed in their rule structure and thus the number and complexity of the operations they required. In particular, we sought to measure the costs of increased WM load and the information manipulation required by the N-back tasks.

Comparing the subjects’ fair wage demands for the 1- and 2-back tasks allowed us to measure the cost of maintaining one more item in working memory (“maintenance”). Comparing the demands for the 2-back and 3-detect tasks, which both require the maintenance of 2 items, allowed us to measure the cost of protecting against interference in the contents of WM (“interference”). In the 3-detect task, subjects had to remember the 2 previous stimuli and compare them to the current stimulus. Detecting a match was simple as long as one recalled whether the previous 2 stimuli matched the current one. In the 2-back task it remained essential to recall the previous 2 stimuli. However, the stimulus from 1 trial ago was never relevant for the trial at hand; all that mattered was the identity of the stimulus 2 trials ago. Because both must be stored, however, it is possible that the stimulus from 1 trial ago was distracting, and if it matched the current stimulus, it may have served as a lure to respond. Identifying and filtering out this distraction may require significant attention and effort. Thus a “lure trial” was any trial where the irrelevant stimulus from 1 trial ago matched the stimulus on the current trial, in the 2-back task. The interference cost in our model captured the cost of these lure trials. This idea is also described in (52).

Stimuli were presented in pseudo-random order such that the use of other WM operations, like WM updating, also differed slightly across rounds. Additionally, forcing 3-5 matches per round allowed us to measure the costs of responding to perceived matches, not responding when matches occur (misses), or responding erroneously (false alarms).

To obtain fair wage ratings for each round in each task, we employed an inversion of the typical Becker-Degroot-Marschak (BDM) auction procedure, in which subjects bid for items with points. In our procedure, subjects are asked to do some cognitive work in exchange for a fair wage. Before each round, subjects were shown a fractal image associated with one of the tasks, and were asked to use a slider to specify their “fair wage” for completing one round of that task. Possible fair wages ranged from 1 to 5 points. They were then shown a random computer offer, also from 1 to 5 points. If the computer offer was above their requested wage, they were given the computer offer for completing one round of the task associated with the fractal. If the computer offer was below their requested wage, they completed the default task for 1 point. All task rounds consisted of 15 trials.

We used the BDM procedure in this work because, via mechanism design, it motivated subjects to report the true subjective value of the effort they expected to expend on each instance of a task. If subjects were effort avoidant and wanted to earn higher wages or not complete effortful tasks at all, they would ask for high wages. If subjects were effort seeking, or at least not effort avoidant, then their fair wages would be low as they should be satisfied with any number of points above the minimum. If one task was substantially more effortful, then our subjects should ask for higher wages on that task so that they would not have to complete that task without proper compensation. In an attempt to prevent subjects from being overly avoidant of making errors, we did not impose an accuracy cutoff for the receipt of points on individual rounds. Further, after the initial practice phase, subjects were not informed of their accuracy each round. However, subjects were aware that if they were inattentive to the task, or their overall accuracy fell below some cutoff, that the task would conclude early and they would receive less compensation (see exclusion criteria below). At the end of the task, subjects’ points were tallied and converted into a monetary bonus.

At the end of the main experiment, subjects completed a basic demographic inventory, the Need For Cognition Scale (NFC; 2), and the Short Almost Perfect Scale (SAPS; 29). They then rated the difficulty of each of the tasks (signaled by its associated fractal) using the same slider that they used to provide their fair wage ratings. Subjects were also able to provide comments on their experiences completing the experiment. Subjects were given one hour and 15 minutes to complete the entire experiment.

### Recruitment and exclusion criteria

The subject pool was limited to Amazon Mechanical Turk workers based in the United States, to ensure English reading comprehension. We limited our recruitment to workers ages 18 and up with at least 100 completed Human Intelligence Tasks, and with at least 85% acceptance rates. We also ensured that subjects had not completed the task before using their Worker ID. To ensure that subjects understood the task and were able to maintain a high level of accuracy, we excluded subjects who did not demonstrate task proficiency or an understanding of the fair wage procedure after the practice phase. We implemented two tests that subjects had to pass to make it into the main experiment. First, subjects had to reach 80% task accuracy on 15 trials of our most difficult task, the 2-back. They had up to 10 rounds to do so. 52 subjects failed to reach this criterion. Following that, subjects had to correctly answer 4 out of 6 questions about the BDM procedure. 76 subjects did not pass this quiz. If subjects passed both those checks, then they proceeded to the main experiment. After these exclusions, 142 subjects started the main experiment.

During the main experiment, subjects’ performance was assessed 3 times (every 8 rounds). If in 8 rounds, subjects missed the response deadline for 4 fair wage ratings or their overall accuracy went below 60%, the task ended early and their data were not used in the final analyses. This eliminated another 42 subjects, resulting in a sample size of 100 subjects total. Subjects were given a 30 second rest between task rounds and no other breaks.

### Model-agnostic analyses

All model-agnostic and model-based analyses were run in MATLAB (53). Subject accuracy was calculated online as a weighted function of correct responses (hits) and correct withholding of responses (correct rejections), where hits were given three times more weight than correct rejections. We chose to emphasize hits over correct rejections in order to encourage participant engagement in the tasks, though subjects were not aware of the exact scoring procedure. In this way, subject accuracy was tracked while they completed the experiment, so that subjects who were not engaging with the task could be removed from the experiment early. Once subjects completed the experiment, we examined their behavior on each task by running ANOVAs on accuracy, response time, fair wages, and difficulty ratings, looking for an effect of task identity. We examined significant main effects of task identity with post-hoc t-tests. We assessed the linear relationships of mean accuracy on each task and mean fair wage for that task across subjects. Additionally, we ran linear analyses of accuracy versus task iteration and overall experimental round, to examine potential learning or fatigue effects on accuracy. We ran these same analyses on fair wage demands to determine whether subjects’ fair wages may have changed with time or task practice. Additionally, we ran a comparison of reaction times on fair wage ratings at the start and end of the experiment.

We scored subjects Short Almost Perfect Scale (SAPS) and Need for Cognition Scale (NFC) responses by summing the numerical values of all their answers, reversing some values as indicated by published scoring guidelines, then dividing by the number of questions answered. We used this normalization to ensure that any questions that were not responded to would not artificially lower questionnaire scores. We excluded questionnaire data from subjects who incorrectly answered one or both of our screener questions (i.e. “Please select ‘Strongly Agree’ for this question”). This type of attention check has been shown to be a reliable way of removing subjects who are randomly responding to questionnaires, especially when administered more than once during an experiment (54).

The mean(std) normalized NFC score was 3.4(0.9) and the mean(std) normalized SAPS score was 4.4(1.3). 1 subject chose not to finish those questionnaires and as such has no NFC or SAPS score.

We correlated these questionnaire scores with each other and with participant age. NFC and SAPS scores were positively correlated (r = 0.24; p < 0.05). There was no relationship between participant age and NFC (NFC/age r = −0.17; p > 0.1) or SAPS score (SAPS/age r = −0.16; p > 0.1). We also regressed NFC and SAPS scores, and their squares, against mean fair wage ratings, average accuracy, and average response time. We used a model selection procedure which trimmed each regression down to an intercept term, and the self-report terms which were necessary for model significance (p < 0.05). If two reduced models were significant, and they included different terms, we selected the model with the lower mean squared error (MSE) in predicting each task variable. We did this to assess both linear and quadratic relationships between individual difference scores and task performance measures.

To build upon the quadratic relationships observed, we also split subjects into tertiles based on their questionnaire scores. Because both scales administered were short-form, many subjects have the same score. Thus after splitting subjects into low, mid-, or high scoring groups based on these scores, the resulting tertiles did not have the same number of subjects in them. Nevertheless, we ran a series of ANOVAs and post-hoc t-tests to examine whether these groups differed in their task accuracy, or fair wages.

### Computational methods and model-based analyses

We use a computational model to quantify the putative cognitive processes used in task completion and their influence on fair wage ratings. We use a process model to decompose each task into the cognitive operations putatively involved in its completion. We fit 84 candidate models to subject data. We fit this many models in an attempt to account for most possible combinations of cost parameters, while also limiting model fitting to those models with high individual parameter recoverability. This number is also elevated by our use of two different functions of how fair wages change with time. We modeled subjects’ fair wage ratings as a dynamic process driven by subject learning or by the changing costs of cognitive effort. The first class of models tests the hypothesis that the total cost associated with each task is learnt through experience with the task and the number of costly components required to complete it (ɑ class of models). The second class tests the hypothesis that cognitive effort costs may themselves change over time, as costly processes become either less costly with practice or more costly as subjects grow fatigued (δ_j_ class of models).

The fair wage ratings for each task were initialized in the model by fitting initial rating parameters for each task and each subject, thus capturing each subjects’ initial ratings with very high fidelity (Supplementary Figure 4). Each subject’s initial fair wage ratings for each task were captured using a free parameter *init_i_*.

rating^0^(task = “1-back”) = *init_1-back_;*

rating^0^(task = “2-back”) = *init_2-back_;*

rating^0^(task = “3-detect”) = *init_3-detect_*

While the inclusion of three extra free parameters to determine initial fair wages may seem unnecessary, correctly capturing each subject’s starting point allows us to fit most accurately how subjects’ fair wage ratings evolve over course of the experiment, as well as how they respond to individual cost components. However, because there are already extra parameters in the δ_!_ class of models (+1 δ_j_ for each cost parameter, so that they can change independently), we did not fit individual *init* parameters to each task in this class of model, to avoid overfitting. After the initial fair wage ratings, the total cost on task round r_k_ of task k was then used to determine the fair wage rating on the next round of that same task (round r_k_ + 1). This round may arise some trials later; we denote trials by t.

We approached cost decomposition with a simple program which was capable of accurately completing each task with the same “cognitive” functions, but switched between rule structures depending on the task at hand. We tallied each operation that the model had to use to complete each task round with 100% accuracy, including how many items had to be maintained in WM, how many times WM storage had to be updated with new information, or how many times there were interfering “lure” stimuli in WM storage. In addition, we tallied the mistakes (misses and false alarms) and button press responses made by each subject in each round. All those components were then scaled by their associated costs (which might change over trials t, and were fit through the modeling), and were summed to produce the total cost incurred on that round of that task. For round of task k:

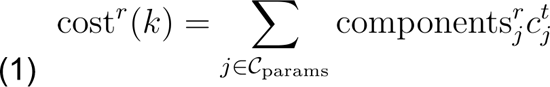

The most complex model included six cost parameters (set *C_params_*): the cost of responding to a perceived match (c_response_), the cost of maintaining information in WM (c_maintenance_), the cost of protecting against interference in the contents of WM (c_interference_), the cost of updating WM with new information (c_update_), the cost of false alarm responding when there was no match (c_fa_), and the cost of missing a match (c_miss_). Other than the interference cost, which was only present in the 2-back task, each cost was fit from ratings of all 3 rated tasks. However, we tested models containing all combinations of 6 different possible costs. All cost parameters were unbounded such that they could be positive, or negative. If any components were perceived to be rewarding, instead of costly, then our model would capture that with a negative cost magnitude. It is important to note that a different choice of process model could result in a different cost structure.

We tested two possible fair wage rating updating mechanisms: a class of model which assumed that the costs themselves changed over trials and that subjects directly reported their experienced costs as they changed, and a class of model which learned the total cost of completing each task following task experience. These updating mechanisms are subtly different, and involve two different free parameters: δ, the scalar with which costs are changed trial-by-trial, and α, the cost learning rate. It should be noted, however, that it is theoretically possible that both mechanisms contribute to cost ratings simultaneously. For simplicity and for robustness of model recovery, we chose to fit these updating mechanisms as separate model classes.

In the cost-changing class of models, δ_j_ (*j* ∈ *C_params_*) is the cost-specific change parameter which captures how costs linearly change over time (trial number t), i.e. with task experience or fatigue:

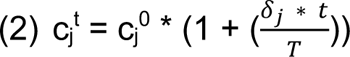

*T* is the total number of trials, and δ can be positive or negative. The flexibility of δ allows the cost of each cognitive operation c_j_ to increase or decrease linearly. Note that because the cost components are shared over tasks, and fatigue is supposed to generally increase with time on task, in this model class each cost is changed according to overall trial number (t), instead of task round number (*r_k_* for task *k*). In this class of models, fair wage ratings on round *r_k_* + 1 are a direct function of the cost parameters and task components involved to complete the previous task round *r = r_k_* (which is equivalent to having a cost learning rate α = 1):

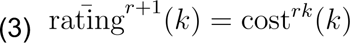

In the cost-learning version of the model, the costs do not change with trial number, as they do in the other class of models, so: c_j_^1^ = c_j_^2^ = … = c_j_^T^. This class of models learns incrementally about the total cost of completing each task. α is the subject-specific cost learning rate which captures how much each subject adjusts their ratings for an individual task based on the most recent round *r = r_k_* of that task:

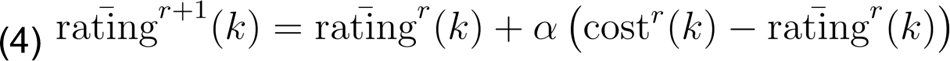

Lastly, we modeled noise in the fair wage rating process with a Gaussian noise process centered on 0 with standard deviation, also a free parameter, and by applying this noise to each fair wage rating independently. This makes the generated rating follow:

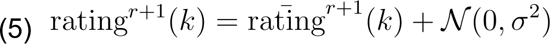

Given the modest number of ratings provided by each subject (32 in total, split amongst 3 tasks), and the overall similarity of ratings between subjects, we fit our models using a hierarchical Bayesian inference (HBI) for computational behavioral modeling (CBM) package (31). Employing a hierarchical parameter estimation procedure allows for similarity across subjects to be leveraged to fit individual parameter values accurately, especially when fitting few individual data points. The package leverages estimations of group parameter means and variances in the individual parameter estimation process. In addition, this package allows for the possibility that not every subject is best fit by one model. Model responsibilities are calculated subject-by-subject such that subjects who are not well-described by a model do not influence the overall parameter probability distributions from that model. In our case, this allows for individual differences in what processes are perceived as costly. If the ratings of some subjects are not affected by a certain cost term, then the group-level estimate of this cost is not driven down by their inclusion in the pool. The Bayesian model fitting procedure constrains the group parameters to have Gaussian distributions, and so, as is common, we transformed the parameter associated with the learning rate using a logistic sigmoid (so it lies between 0 and 1) and the parameter associated with the rating noise using an exponential, so that it is positive (with a log normal distribution).

To validate the winning models further by assessing their ability to produce the behavioral effects of interest, we simulated fair wage ratings using each of the winning models. We then compared these model simulations to real subject behavior via visual inspection, and by computing mean r-squared values for each model. Because stochasticity is one feature of model behavior (via the standard deviation parameter σ), we simulated each subject’s data using their fit parameter values 10 different times to control for the stochasticity of these simulations. Each time, we correlated the true fair wage ratings of all subjects with the set of simulated fair wages, and then squared the r-value obtained. We ran this over 1000 iterations, and then took the average r-squared value to produce a mean r-squared value for each model. This was then used to validate that the models could reproduce subject behavior.

In the CBM toolbox, the group-level mean for each parameter is calculated separately for each model. This allows group-level cost parameter magnitudes to be compared within-model, but not across-model. In order to compare the magnitudes of the cost parameters across all our models, we constructed posterior probability distributions over the magnitude of each cost. We used parameter estimates from every subject and every model, weighing the contribution of each subject *s* and model *m* by their fit responsibility ρ:

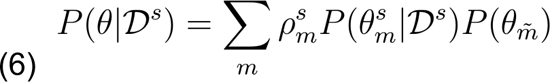

where *P(θ_m^_* is the group-level prior distribution over the costs not in model *m*. This prior is a weighted average over the group-level parameter distributions derived from each model, where the weights are again derived from the model responsibilities *ρ^s^_m_*. We assumed that these prior distributions were Gaussian within-model, then averaged them across models to produce non-Gaussian mixture models of across-model priors. Here, the responsibility *ρ^s^_m_* reflects the degree to which each subject’s fair wage data were explained by that model.

Using equation 6, we constructed a 4D distribution over the four cost parameters included in models with at least 1% fit model frequency. We summed over the 4D joint distribution to produce the marginal distributions of each cost. Additionally, we subdivided our subjects into tertiles based on self-report scores (NFC and SAPS), and calculated the degree to which the posterior parameter distributions overlapped across these score groups *g*:

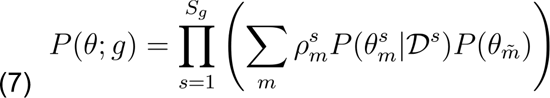

where subjects 1 through *S_g_* belong to the group of interest.

We obtained the means and standard deviations of the marginal posterior distributions over individual cost magnitudes. In this way, we assessed the degree to which the cost magnitudes were separable within- and across-subjects, and across models which did not share all the same parameters.

We confirmed the validity of our models and model fitting procedure by running a generate and recover procedure. For each model, we simulated a data set of 30-100 subjects with known parameter values. We used trial-by-trial cost components taken directly from subject behavior to ensure that real responses, including errors, and task characteristics were compatible with our modeling procedure. To determine which models were sufficiently robust in parameter recovery, we ran a generate and recover of all 126 possible models (combining different costs and using an α or δ update mechanism). In this way, we selected 84 models to test that showed reliable parameter recovery and minimal cost parameter tradeoff. We wanted to test a broad array of models since we had limited a priori knowledge of which cost components would drive fair wages, or what form cost updates would take. At the same time, we wanted to fit real subjects’ data only with models that had recoverable free parameters and minimal tradeoff between costs, despite possible correlations of cost components, as individual differences were of particular interest.

Supplementary Figure 4 shows the results of this generate and recover procedure for one example model, which includes update, maintenance, and false alarm costs (N = 50 simulated subjects). All fit and real parameters were highly correlated ( r = 0.84, p < 0.001; r = 0.94, p < 0.001; update costs r = 0.55, p < 0.001; maintenance costs r = 0.66, p < 0.001; false alarm costs r = 0.78, p < 0.001; init_1-back_ r = 0.88, p < 0.001; init_2-back_ r = 0.96, p < 0.001; init_3-detect_ r = 0.92, p < 0.001). This indicates that our models supported the reliable recovery of individual parameters, despite the modest number of trials that were fit per subject.

## Acknowledgements

We thank our subjects on Amazon Mechanical Turk for their participation in this experiment, as well as our earliest pilot subjects, the members of Arbeitsgruppe Peter Dayan (AGPD) at the Max Planck Institute for Biological Cybernetics. We also thank the members of AGPD for many helpful comments on the initial design of the experiment, and specifically thank Dr. Franziska Broeker for her assistance in establishing the online data collection pipeline, and Drs. Andrew Webb and Aenne Brielmann for thoroughly reviewing the modeling and analysis code for this project. We thank Dr. Martin Breidt, Dr. Holger Dinkel, Mihai Vintiloiu, and the entire IT Core Facility at the Max Planck Campus in Tuebingen for their technological and research administrative support. We thank Finn van Krieken for his assistance with front-end online experiment development. We also thank Dagmar Maier for administrative support.

## SUPPLEMENTARY INFORMATION

### SUPPLEMENTARY RESULTS

Because the 2-back was associated with the most WM updating, most errors, and most interference in WM, many of the cognitive components of task completion that came out of our process model were highly correlated. For example, across all subjects & task rounds, updates, maintenance, lures, and false alarms were all correlated (interference vs. false alarms r = 0.48, p < 0.001; interference vs. maintenance r = 0.53, p < 0.001; maintenance vs. false alarms r = 0.31, p < 0.001; updates vs. maintenance r = 0.86; updates vs. false alarms r = 0.43; updates vs. interference r = 0.62). In addition, when we ran within-subject correlations of these components across rounds, they were significantly correlated within 91% (maintenance vs. interference), 44% (maintenance vs. false alarms), 74% (interference vs. false alarms), 100% (updates vs. maintenance), 66% (updates vs. false alarms), and 90% (updates vs. interference) of subjects. One consequence of this may be that the cost parameter values associated with these components may trade off with one another in model fitting, artificially raising or lowering each other. In addition, model selection may have been impacted, resulting in a low number of subjects who were best fit by models including multiple costs. Because most subjects’ data are best captured by a model including only one cost of cognitive effort, one might wonder whether the cost parameters obtained from our models are capturing one cost only, but incorrectly assigning them to three different components due to the relatedness of the components. To compare cost parameter magnitudes across models including only one or two cost parameters each and to ensure their separability, we constructed posterior distributions over parameter values using the outputs of the CBM toolbox(31). We also examined whether these parameter values traded off during model fitting by examining their covariances, which are derived from the inverse Hessian of the search gradient within the multidimensional parameter space.

The most frequent model with multiple cost parameters contained update, maintenance, and interference costs. The covariances between these parameters, which is influenced both by their empirical covariances, and their covariance during parameter fitting, were all within an acceptable range. The covariance between the update and maintenance costs was largest, at −0.22. Between the update and interference costs, the covariance was 0.0492. Between the interference and maintenance costs, it was −0.0763.

We verified in a pre-model fitting generate and recover procedure that individual cost parameters were being accurately fit even in models with multiple costs (Supplementary Figure 4). In addition, there was no evidence that these single cost parameters somehow capture just one underlying component, rather than 3 separate ones, as the posterior distributions over their magnitudes are mostly non-overlapping on the group level (Figure 3B). While the update and interference costs are of similar magnitudes, and therefore overlapping, the negligible covariance between update and interference costs suggests that they did not trade off in model fitting.

## SUPPLEMENTARY FIGURES

**Supplementary Figure 1.**
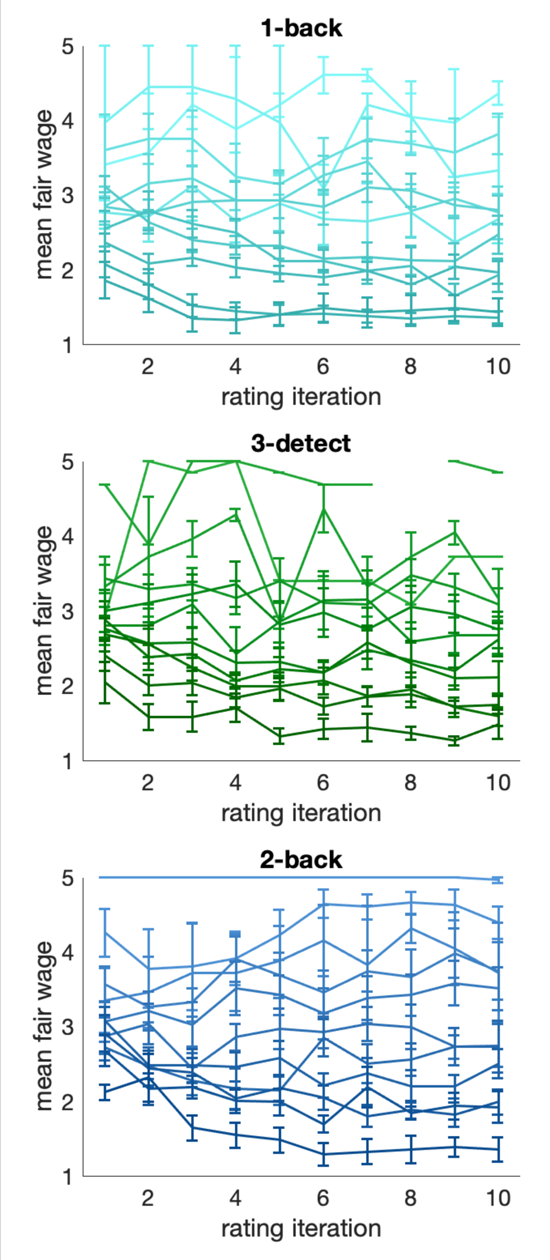
Mean fair wage ratings across task rating iteration. All subjects completed 10-11 ratings of each task, but only between 1 and 11 rounds per task. Here we plot the mean fair wage for the subjects who completed 1 to 11 iterations of each task, grouped by the total number of iterations they completed. Subjects who completed more task iterations are plotted in darker colors. This illustrates the diversity in fair wage ratings for each task across subjects, as well as the stability of the ratings subjects gave to each task. In addition, it shows that, due to the design of our task, subjects who asked for high fair wages on one of the tasks did indeed complete fewer iterations of that task. Error bars are drawn with standard error of the mean.

**Supplementary Figure 2.**
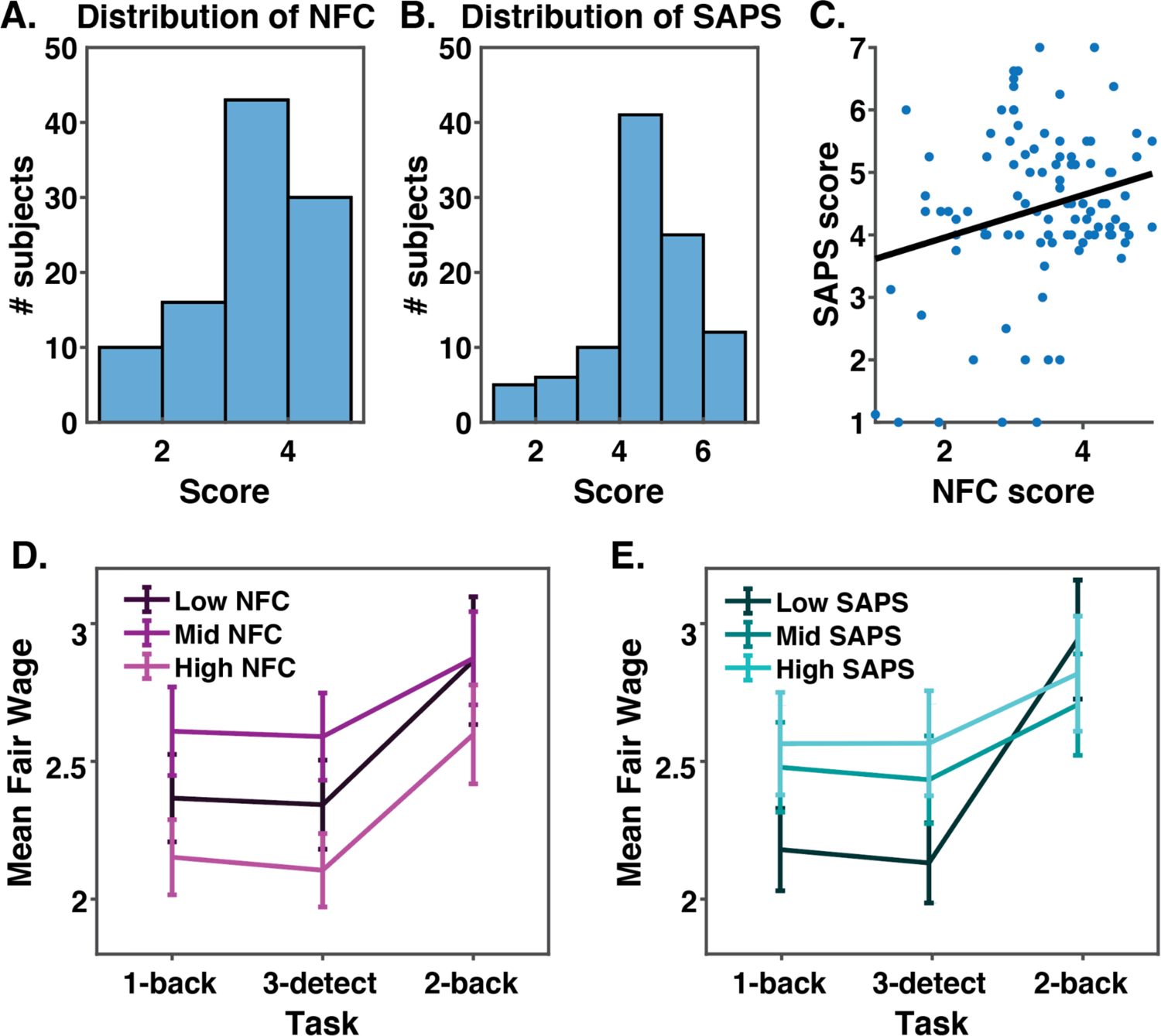
Self-report scores and their relationships to mean fair wages. **A.** Distribution of Need for Cognition (NFC) scores within the experimental sample. Scores have been normalized by the number of questions answered such as not to lower the mean of the distribution artificially. The distribution of NFC scores in our sample is right-skewed compared to the typical distribution of NFC scores. However, this is typical of subjects on MTurk (Berinsky, Huber & Lenz, 2012). **B.** Distribution of Short Almost Perfect Scale (SAPS) scores. Scores have been normalized by the number of questions answered such as not to artificially lower the mean of the distribution. The distribution of SAPS scores in our sample is typical of both in-person samples (Rice, Richardson, & Tueller, 2013) and other samples on MTurk, including one sample of 400 subjects (Stricker, Flett, Hewitt, & Pietrowsky). **C**. NFC scores versus SAPS scores. NFC and SAPS scores were positively correlated (r = 0.24; p < 0.01). **D.** Mean fair wage rating on the 1-back, 3-detect, and 2-back tasks by tertile split NFC groups. Error bars were drawn using the standard error of the mean (SEM). There was a significant quadratic relationship of NFC and mean fair wage ratings (β = −0.03). Post-hoc t-tests confirmed that the significant quadratic effect of NFC was only driven by mid NFC subjects having significantly higher fair wage ratings than high NFC subjects (p < 0.01). **E.** Mean fair wage rating on the 1-back, 3-detect, and 2-back tasks by tertile split SAPS groups. Error bars were drawn using the SEM. A 3-way ANOVA, revealed no effect of SAPS group (F = 2.2, p > 0.05) or of the interaction of SAPS group and task identity (F = 1.5, p > 0.05) on fair wages.

**Supplementary Figure 3.**
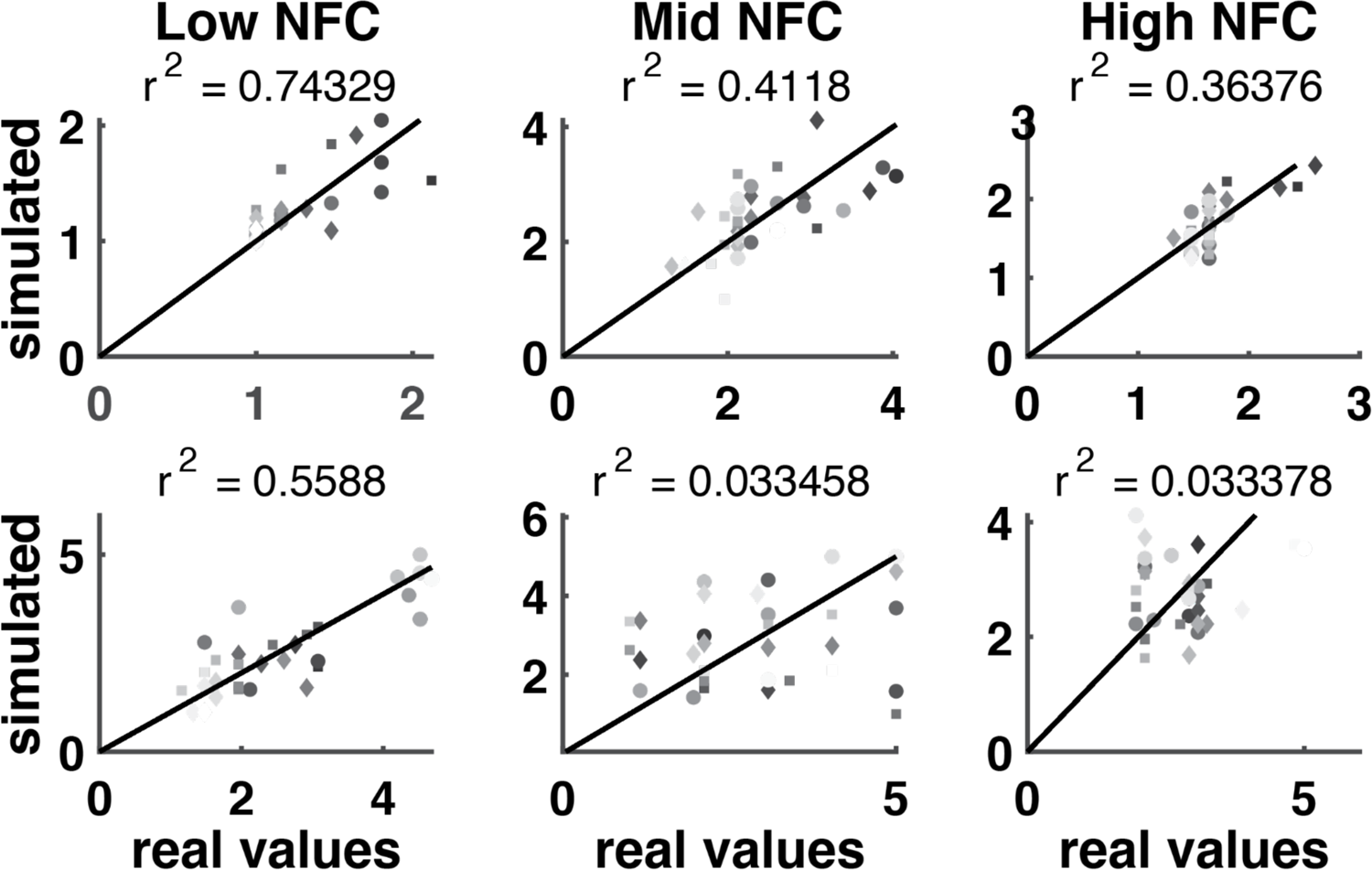
Real fair wage values versus simulated fair wage values. For each subject, we simulated their data using the model which took the highest responsibility for their data, and their fit parameter values. Here we have selected 2 random subjects from each NFC tertile (left: low NFC, middle: middle NFC, right: high NFC) and plotted their real and fit fair wage values. The title of each plot is the mean r-squared value after 100 simulations with the subject’s best fit model and best fit parameter values. Markers are shaded such that later trials are displayed in darker colors, and the shape of the marker indicates which task was rated (squares are 1-back ratings, circles are 3-detect ratings, and diamonds are 2-back ratings).

**Supplementary Figure 4.**
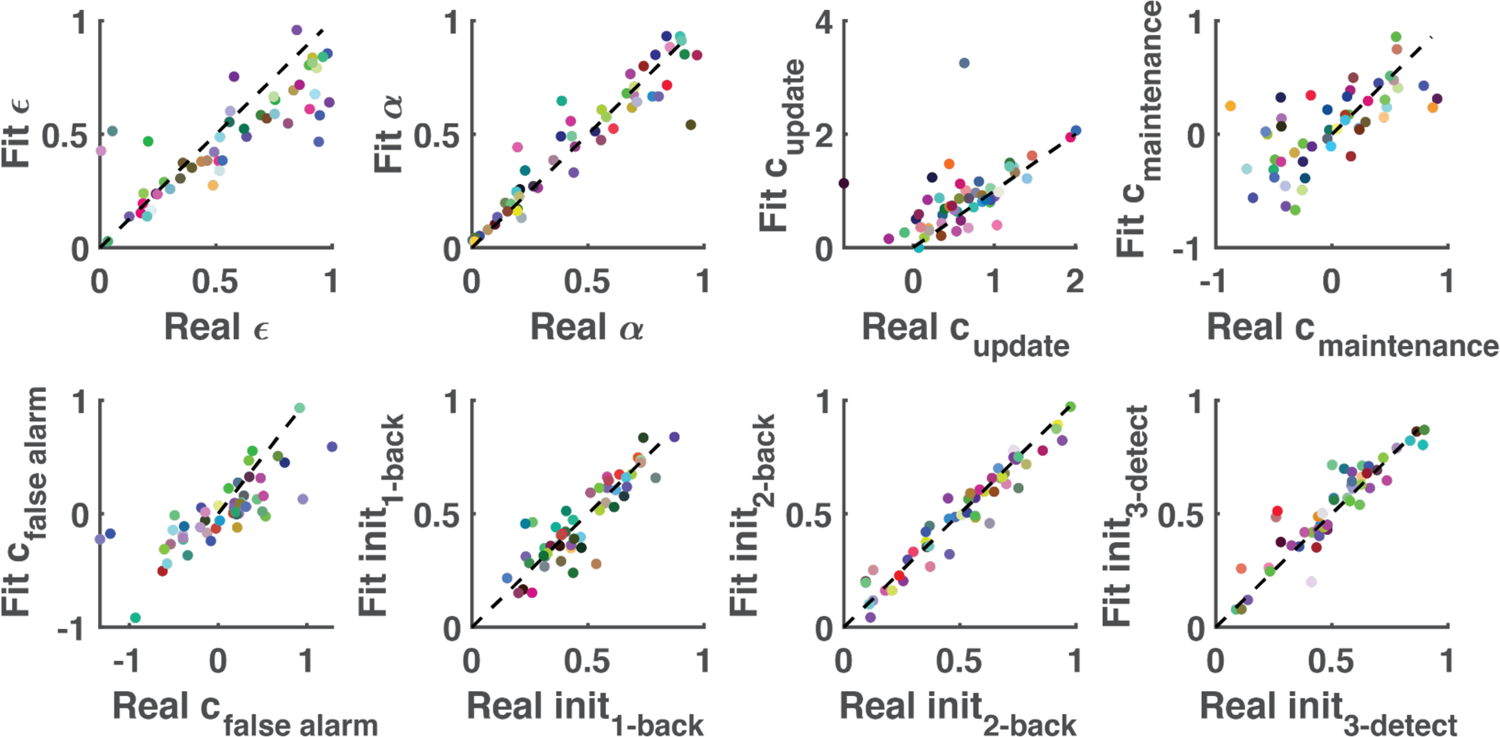
The results of a generate and recover procedure on a model including update, maintenance, and false alarm (FA) costs. A dataset of simulated subjects was produced with random parameter values (constrained by the bounds of those parameters), and then fit with the same procedure as real subject data. Here we show the fits for 50 subjects out of 100, where each subject’s fits are plotted in a unique color. The identity line is overlaid on each subplot in black. Comparing the fit parameter values to the real values reveals the high fidelity of the model fitting procedure. Models were fitted using the Computational Behavioral Modeling (cbm) toolbox of Piray et al (2019). All candidate models were visually inspected and verified as recoverable to avoid fitting models with parameter tradeoffs. Only models with parameter recoverability were fit to real subject data.

